# Rapid DNA Re-Identification for Cell Line Authentication and Forensics

**DOI:** 10.1101/132381

**Authors:** Sophie Zaaijer, Assaf Gordon, Daniel Speyer, Robert Piccone, Yaniv Erlich

## Abstract

DNA re-identification is used for a broad range of applications, ranging from cell line authentication to crime scene sample identification. However, current re-identification schemes suffer from high latency. Here, we describe a rapid, inexpensive, and portable strategy to re-identify human DNA called MinION sketching. Using data from Oxford Nanopore Technologies’ sequencer, MinION sketching requires only 3min of sequencing and ∼91 random SNPs to identify a sample, enabling near real-time applications of DNA re-identification. This method capitalizes on the vastly growing availability of genomic reference data for individuals and cancer cell lines. Hands-on preparation of the samples can be reduced to <1 hour. This empowers the application of MinION sketching in research settings for routine cell line authentication or in forensics.

Software is available at <u>https://github.com/TeamErlich/personal-identification-pipeline</u>

## Background

DNA is a powerful biometric identifier. With the exception of monozygotic twins, DNA profiles are unique to each individual on Earth (Kayser & de Knijff 2011; Bieber et al., 2006; Gymrek et al., 2013). The ability to re-identify DNA has multiple applications in a broad range of disciplines. In research settings, re-identification is employed to authenticate cell lines by matching their DNA to validated genomic profiles (NIH 2016; AMS 2015). In clinical genetics, the American College of Medical Genetics recommends using companion DNA genotyping tests to track sample identity to avoid sample mix-ups during clinical whole genome/exome sequencing (Green et al., 2013). In forensics, DNA identification has become one of the most common techniques to identify crime scene samples, casualties of mass disasters, and victims of human trafficking (US Deptartment of State, 2014).

Despite this wide range of applications, current DNA identification methods suffer from high latency and low portability. Numerous recent reports have highlighted the high prevalence of mislabeled cell lines that result in irreproducible research and squandered scientific funding (Almeida et al. 2016; Chatterjeem 2007; Dolgin & Elie 2016; Capes-Davis & ICLAC 2016; Nardone, 2007; Simeon-Dubach et al., 2016). To mitigate this issue, the NIH and various journals require researchers to authenticate cell lines by matching their DNA profiles to validated signatures (NIH, 2016; AMS, 2015). Currently, the most common DNA identification strategy genotypes a small set of autosomal polymorphic short tandem repeats (STRs) (Smith et al., 2012; Capes-davis et al., 2010; Reid Y et al., 2013; Masters et al., 2001; ATCC 2011). But this technique requires time consuming PCR-based steps and specialized capillary electrophoresis machines. In forensics, the state-of-the-art DNA identification platforms (e.g. DNAscan or RapidHIT200) take about 90 minutes to process a DNA sample, weigh over 50 kilograms, have a capital cost of more than $250,000 and require about $300 to process a sample (Hennessy, 2013). While the American Type Culture Collection (ATCC) offers an STR-based cell identification service for $195 per cell line, the overall procedure requires shipping consumables and samples back-and-forth and takes two weeks to complete. A recent survey reported that the delay in research is one of the primary reasons researchers avoid cell line authentication (Almeida et al., 2016). Previous studies have considered using SNPs for re-identification but are yet to address the latency issue. Indeed, a carefully selected panel of ∼50 SNPs confers a re-identification power similar to that provided by the 13 STR markers used in forensics (Sanchez et al., 2006; Yu et al., 2015). Nonetheless, genotyping these SNPs requires PCR amplification genotyping technologies such as Illumina sequencing, Sanger sequencing, or SNP arrays, all of which have relatively long processing times of usually over a day, and suffer from the absence of portability and instant accessibility.

Here, we report a portable, rapid, robust and inexpensive strategy for SNP-based human DNA re-identification using a MinION sequencer (produced by Oxford Nanopore Technologies, ONT), a cheap and portable DNA sequencer that weights only 100grams and can be plugged into a laptop computer. This device can be adopted easily in a standard laboratory. Our strategy, termed ‘MinION sketching’, exploits real-time data generation by sequentially analyzing extremely low coverage shotgun-sequencing data from a sample of interest and comparing observed variants to a reference database of common SNPs (Figure 1). We specifically sought a strategy that does not require PCR to eliminate the latency introduced by DNA amplification and to increase portability and miniaturization. However, this poses two technical challenges. First, MinION sequencing exhibits a high error rate of 5-15% (Ip et al., 2015), which is two orders of magnitude beyond the expected differences between any two individuals. Second, MinION sketching produces shotgun-sequencing data that only covers a fraction of the human genome due to the limited capacity of a MinION flow-cell. As such, the extremely low coverage dictates that each locus is covered by up to one sequence read, which nullifies the ability to enhance the signal by integrating multiple reads or observing both alleles at heterozygous loci. Taken together, these challenges translate to a noisy identification task where the available genotype data only provide a mere sketch of the actual genomic data.

**Main figure 1.**
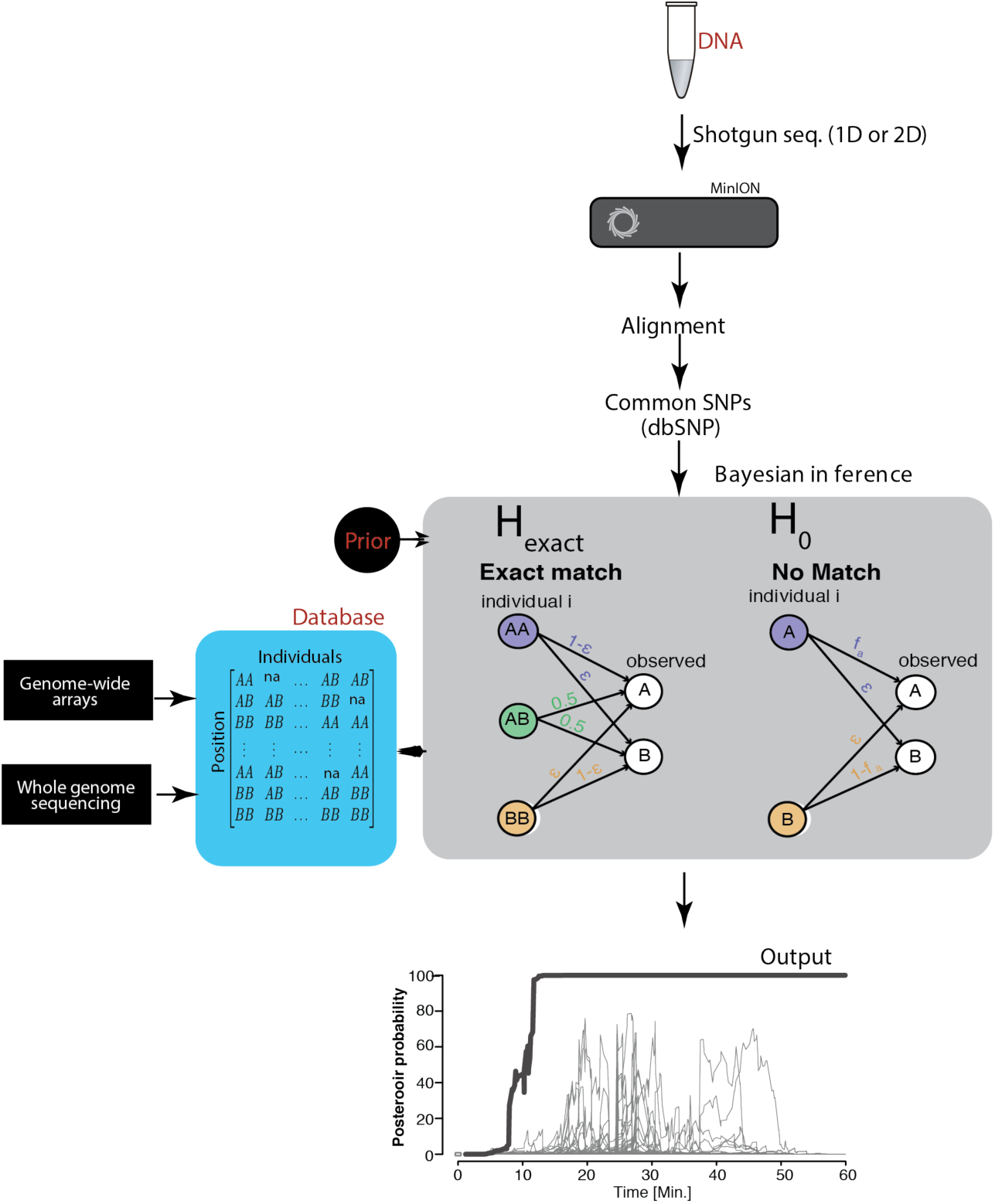
Schematic overview of MinION sketching. A DNA sample is prepared for shotgun sequencing. Libraries are prepared either for 1D or 2D MinION sequencing (e.g. 2D is with hairpin, 1D is without hairpin). Variants observed in aligned MinION reads are only selected if they coincide with known polymorphic loci while others are treated as errors. These SNPs are compared to a candidate reference database comprised of samples genotyped with WGS or sparse genome-wide arrays (∼500K SNPs per candidate file). A Bayesian framework computes the posterior probability that the sample matches an individual in the database by accounting for the sequencing error rate (*ϵ*). This results in an output plot where the posterior probability is visualized as a function of time and the number of SNPs used in the computation.

To address these challenges, we developed a Bayesian algorithm that computes a posterior probability that the sketch matches an entry in the reference database (H_exact_), or has no match to the data data, taking into account each marker’s allele frequency, and the prior probability that a sample matches an entry in the reference database. The Bayesian approach sequentially updates the posterior probability with every new marker that is observed until a match is found. Collectively, our method can identify a sample, without PCR amplification, yet with very high probability despite the low coverage and the high error rate of the MinION.

## Results

We sought to test our strategy using a large-scale reference database and in various technical scenarios in order to benchmark our re-identification method for real-life scenarios. To this end, we first constructed a large-scale reference database of genomic datasets to stress the specificity of our method. This reference database comes from the DNA. Land project (Erlich, 2015) and contains 31,000 genome-wide genotyping array files of individuals tested by Direct-to-Consumer companies such as 23andMe, AncestryDNA, and FamilyTreeDNA (Figure 2A). Next, we ran MinION sketching on four DNA samples in various technical scenarios (Table supplement 1). These scenarios included either extracting the DNA from a spit kit or tissue culture, testing either the R7 chemistry or the newer R9 chemistry, and re-identifying samples that were derived from different ethnic backgrounds. The genetic reference file for each of these samples was included in our database.

We found that the MinION sketching procedure re-identified human DNA with high accuracy after minutes of operation. After only 13 minutes of sketching using the R7 chemistry, the Bayesian algorithm re-identified the NA12890 sample (a female CEU individual from the HapMap project) with a posterior probability greater than 99.9%. Despite the high error rate of this relatively old chemistry and the low coverage, the algorithm needed only 195 bi-allelic variants to re-identify the sample (Figure supplement 1, Table supplement 2), only ∼2 times above the theoretical expectation for re-identifying a person by fingerprinting random markers (Lin et al., 2004). To further test the robustness of our method, we re-sketched NA12890’s sequencing data against reference files for her first-degree relative (NA12877) and second-degree relative (NA12879). Importantly, no exact-matching probability was observed, highlighting the specificity of our method (Figure supplement 1). Next, we repeated the R7 chemistry experiment with another sample of a mixed Ashkenazi-Uzbeki male (YE001). Again, we were able to re-identify this person within 13min and 110 SNPs (Figure 2B, Table supplement 2), further showing that the method produces consistent results across ethnic origins. None of the other 31,000 individuals reached to this level of re-identification (Figure 2B). Finally, we wondered about the impact of the prior probability on identifying individuals. To this end, we tested various prior probabilities of identifying the YE001 sketch. We found that the initial selection of the prior probability had no effect on the matching ability and only slightly increased the time required to achieve a high-confidence match. Even with a prior probability that considers a database around a million times bigger than the world’s population (10^-15^), the posterior probability reached 99.9% with only 25 minutes of sketching YE001 (Figure supplement 2).

Moving to the new R9 chemistry provided even faster re-identification results. We sketched samples of a Northern European female (SZ001) and a Northern-Italian-Ashkenazi male (JP001) using the R9 chemistry. We were able to re-identify these two samples using only 98-134 SNPs and the fastest identification required less then 5 minutes of MinION sketching (Figure 2C, 2D, Table supplement 3). Again, none of the other 31,000 individuals in our database were matched to SZ001 or JP001 using this strategy. The rapid re-identification seems intimately linked to the increased speed of DNA passing through the pore with the R9 chemistry versus the R7 chemistry (250bases/sec *vs* 70bases/sec). These results suggest that further developments in speeding up the DNA reading time can further reduce the re-identification time.

**Main figure 2.**
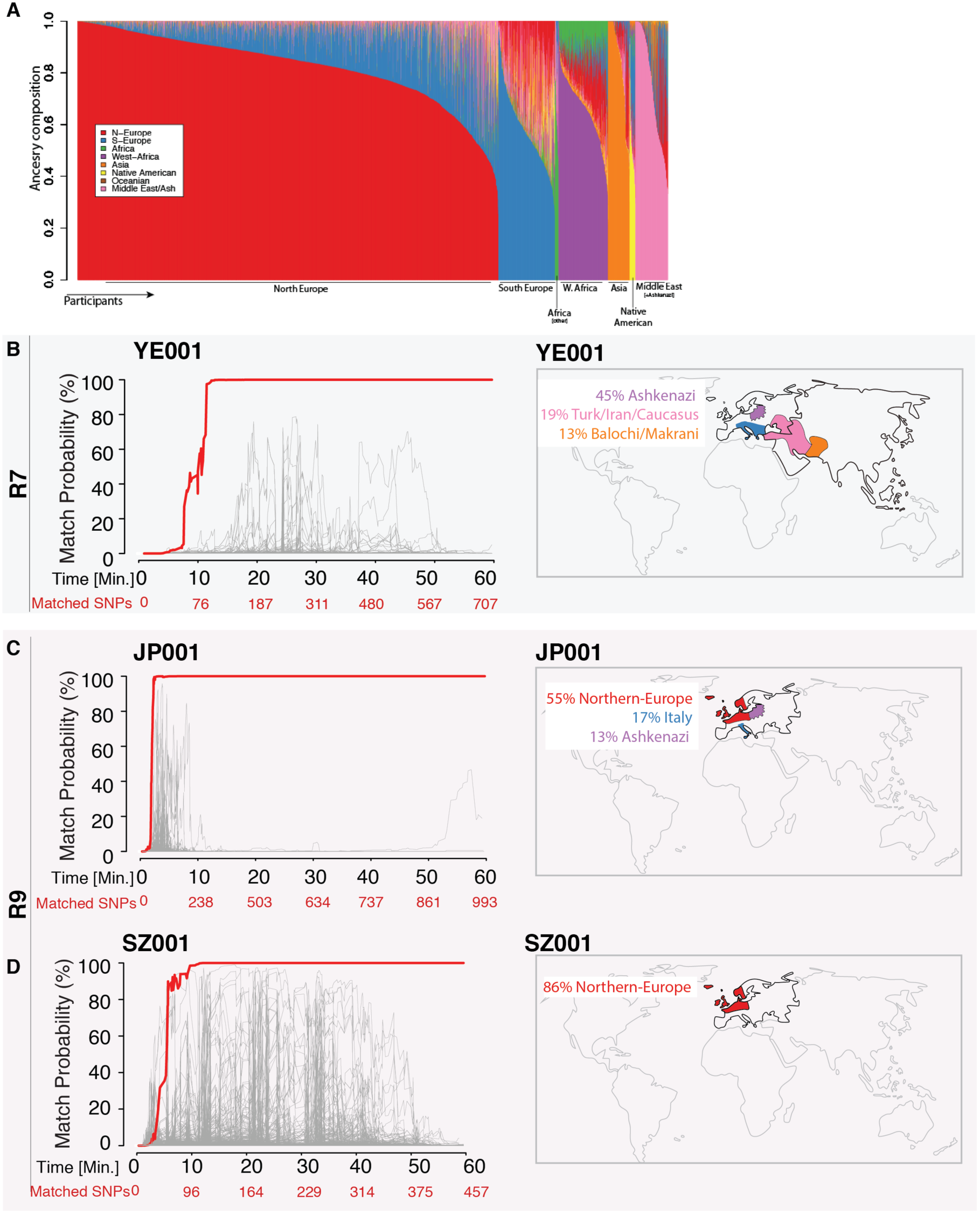
Re-identification of DNA samples. **A)** A Frappe plot showing the population structure of the database with a collection of 31,000 DTC genome-wide arrays. B-D The match probability is inferred by comparing a MinION sketch to their reference file as a function of the MinION sketching time (red line). The prior probability for a match was set to 10^-5^. Matched SNPs (bottom x-axis) denote the number of SNPs used in the posterior computation by the Bayesian algorithm. The match probabilities are inferred by comparing the MinION sketches to a database with 31,000 DTC genome-wide arrays (including the matched individuals). **Right**: Ancestral background is the corresponding individuals; only ancestry predictions of >10% are indicated. (B) The DNA sample was collected from an Ashkenazi-Mizrahi male (YE001) and sequenced using R7 chemistry. (C) Sample was collected from a female North-European (SZ001) and sequenced using R9 chemistry. (D) Sample was collected from a male North European-Italian-Ashkenazi individual (JP001) and sequenced using R9 chemistry.

Next, we explored the applicability of MinION sketching for cancer cell line authentication, a longstanding issue in the research community. To address this, we compiled a collection of genome-wide arrays of 1099 cancer cell lines from the Cancer Cell Line Encyclopedia (CCLE) (Yu et al., 2015; Barretina et al., 2012). These reference files were generated by SNP arrays and contain ∼700K SNP genotypes for each cell line. We then used MinION sketching and the R9 chemistry to authenticate THP1, a monocytic leukemia strain. To show that more than one sample can be authenticated at the same time, we barcoded the THP1 sample and combined it to an additional barcoded human sample. From the barcoded THP1 reads generated in ∼3min of sequencing, the sketching procedure leveraged 91 SNPs to authenticate the THP1 cell-line with a posterior probability of 99.9%. None of the other 1098 CCLE reference files reached a probability of 99.9% or even exceeded 10% match probability (Figure 3A, Table supplement 4).

Next, we wondered about a severe cell line contamination with cells of another origin. Cell line cross-contamination is caused mostly by overgrowth from secondary cell lines with a substantially shorter generation time (Capes-davis et al., 2010). Under the current ASN-0002 standard, a cell line is considered authentic when the STR profile matches to >80% of the corresponding reference panel (Reid et al., 2013; Masters et al., 2001; ATCC, 2011). If the match is <56% it is considered unrelated or contaminated (Reid et al., 2013). To this end, we re-analyzed the data from the THP1 experiment but without resolving the barcodes, essentially reflecting 50% contamination. The algorithm correctly showed a 0% match probability to the THP1 reference file or any other cell line in the database (Figure 3B). We further explored the effect of the faction of contamination on matching the THP1 reference file. By sampling from the above data in different proportions, we found that the algorithm correctly rejects a match for contamination levels above 20% (Figure supplement 3). This shows the that the algorithm will reject authenticating the cell line when there is contamination of over >20%, complying with the ASN-0002 requirements (ATCC, 2011).

**Main figure 3.**
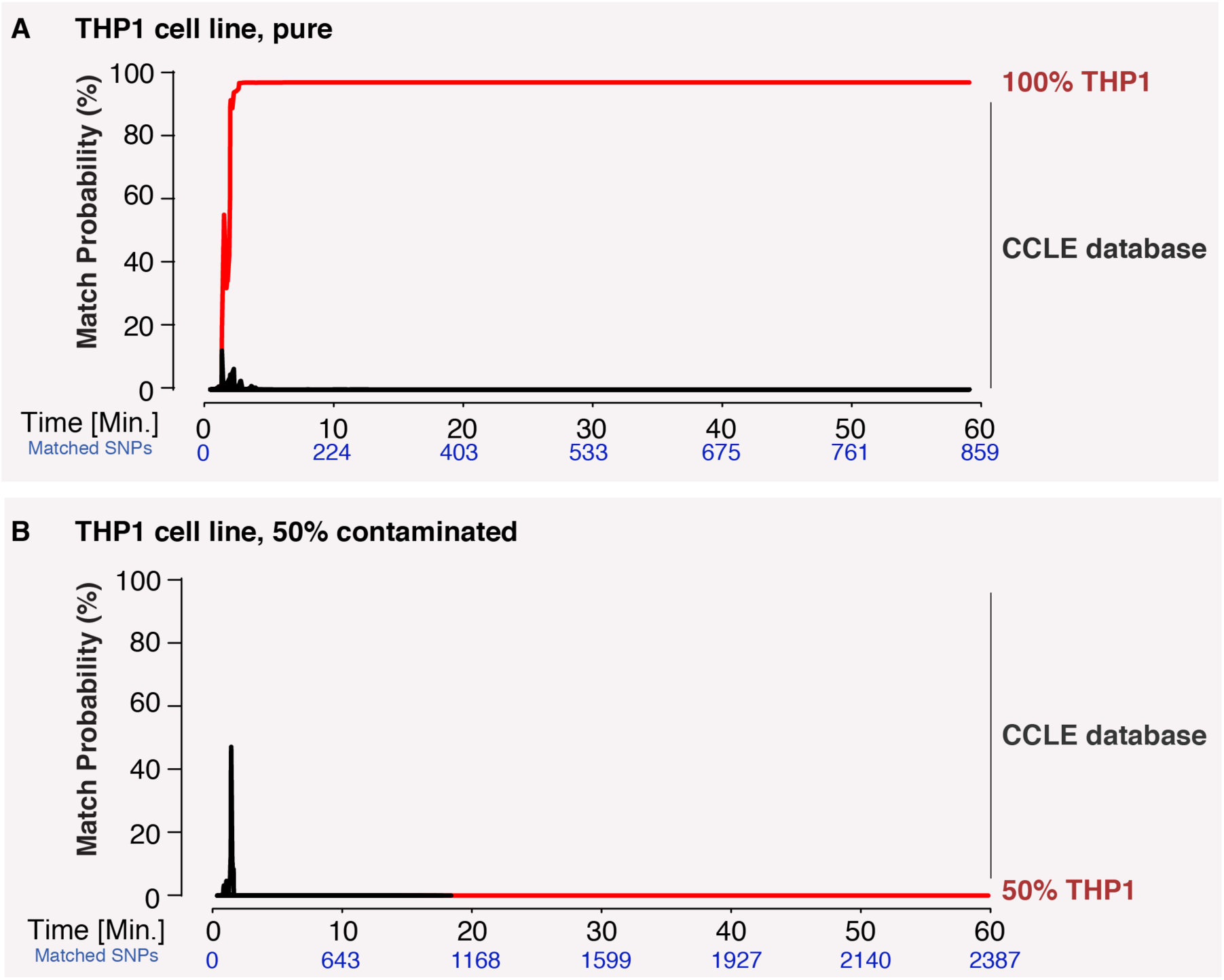
Main figure 3 Cell line authentication. Barcoded DNA from the THP1 cell line is mixed 1:1 with a random barcoded sample. Analysis of only the THP1 reads was used to infer ‘pure’ matches, while analyses of the mixture were used to characterize the efficiency of matching using contaminated samples. The match probability is inferred by comparing a MinION sketch to 1099 reference files that are part of the cancer cell line encyclopedia (CCLE) generated by the Broad Institute (grey). (A) The posterior probability for an exact match between the MinION sketch of the ‘pure’ cell line THP1 (considering a single barcode) and the reference file generated by the CCLE (red is THP1 reference file, other strains are depicted in grey) (B) The posterior probability that the contaminated (50%) mixed sample matched THP1 as a function of the sketching time.

Lastly, we aimed to explore a sample preparation strategy that requires minimal hands-on time. To this end, we utilized a simple protocol to extract DNA using the rapid transposase-mediated fragmentation and adaptor ligation kit provided by ONT. This method generates 1D reads, where only one of the two strands passes through the nanopore, resulting in reads with a higher error rate (Table supplement 5). The advantage of this method is the speed and convenience of the preparation protocol. In only 55 minutes, we were able to extract DNA and produce a ready-to-sequence library (Figure 4A). The increased error rate resulted in the requirement of more SNPs to reach the re-identification threshold. In our experiment, the rapid sample preparation required 239 SNPs after 2.3hrs of sequencing to identify the THP1 cell-line with >99.9% probability (Figure 4B). As such, cell line authentication still can be completed with the same level of accuracy, in one afternoon and using only minimal hands-on time by the researcher.

**Main Figure 4.**
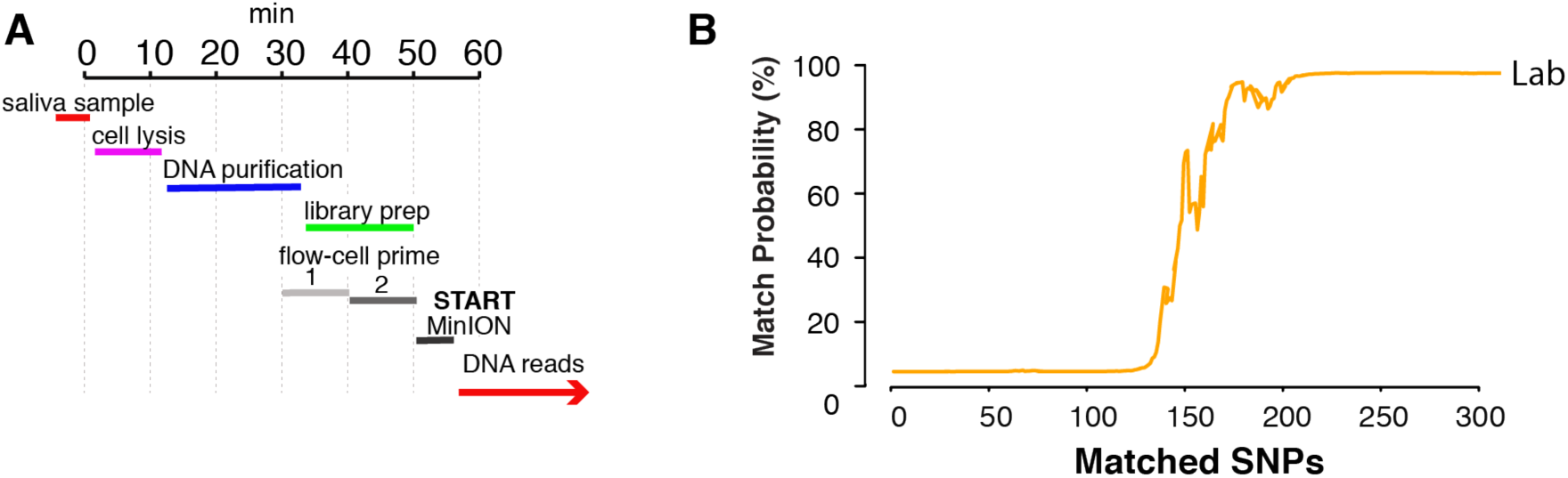
Rapid library preparation. A) Schematic of the steps from sample to MinION sketch. The current method requires ∼55 min until the MinION starts to generate reads. B) The match probability is inferred by comparing a MinION sketch generated by transposase mediated adaptor ligation (the rapid kit) to their reference file as a function of the MinION sketching time (red line). The prior probability for a match was set to 10^-5^. The rapid library protocol was tested in the lab. The MinION sketch generated from sample SZ001. The library was prepared in 55 minutes in the laboratory. After 2.3 hours of sequencing and 239 informative SNPs, the posterior match probability exceeded 99.9%.

## Discussion

Our results show the power of MinION sketching for re-identification of human samples, which can be useful for forensic applications, tracing samples in clinical genetics, and authenticating cell-lines in basic research. Based on only 3-13min of sequencing and 91-250 informative SNPs, MinION sketching can infer the identity of an anonymous sample, and does so robustly, independent of database size and sample ethnicity. MinION sketching is a unique addition to current state-of-the-art re-identification methodologies, because of a number of properties. First, MinION sketching is done using a portable DNA sequencer that can be used in remote locations and therefore reduces the latency of sample transport and sample re-identification speed. Second, by using shot-gun sequencing and intersecting it with the sparse candidate reference file (500K) MinION sketching omits dropouts of informative markers due to sample degradation (Sanchez et al. 2006). Third, the relatively high level of indels in MinION reads nullifies the potential to use STR length polymorphisms for re-identification of DNA samples. Yet, MinION sketching based on SNP-based identification meets the ASN-0002 requirements (ATCC, 2011) for cell line authentication.

Full integration of MinION sketching in forensic settings would require a systematic change of existing standards that rely on STR analysis. Short-term SNP-based re-identification can be applied for crucial identification challenges at mass disasters where new reference files and re-identification are required rapidly. MinION sketching is fully compatible with whole genome sequencing and genome-wide genotyping arrays. Unlike STR profiles, these datasets are much more common in clinical and research settings thus enabling researchers to leverage existing resources for cell line or clinical sample authentication (Barretina et al., 2012). In addition, millions of people have access to genotyping arrays from Direct-to-Consumer (DTC) companies, rendering our method compatible with this type of data as well. Common DTC genotyping datasets can be generated in a highly cost-effective manner (low hundreds of dollars per sample) and within the same price range as the generation of forensic profiles such as the CODIS or ENFSI sets.

We show that cell line authentication can be achieved in the lab in one afternoon, either using a hands-on or hands-off method and be compliant with the ASN-002 standard. In particular, we offered two methods for authentication: the first method involves a hands-on 3hr preparation protocol, but after only ∼3min of sequencing we were able to identify the THP1 cell-line out of 1099 other cancer cell lines with a posterior probability of 99.9%. The second method requires 55mins for the DNA extraction and transposase-mediated adapter ligation and 2.3hrs of sequencing. Both methods take far less time than the two-week process of the American Type Culture Collection. As recent updates to the ONT chemistry (R9.4) have improved sequencing rates to 450 bases/sec, MinION sequencers will likely provide sufficient data for re-identification of a sample in around 1 minute of sequencing. Moreover, multiplexing 12 DNA samples in one run will reduce the cost to a little over $100, which is substantially lower then the ATCC STR-typing service or forensic kits.

As major authentication challenges plague research fields that work with a multitude of plant and mice strains (Petkov et al., 2004; Nitzki et al., 2007; Anastasio et al., 2011; Didion et al., 2014), our work could potentially benefit authenticating samples in remote locations that requires information rapidly and on-site.

The MinION sketches offer a range of capabilities desirable in forensics such as extreme portability and online identification. Early access users have generated MinION sequencing data in unconventional places, including rural Africa (Quick et al., 2016), hotel rooms, and classrooms (Zaaijer & Erlich, 2016). We therefore envision that our strategy can set the basis for near real-time DNA surveillance for forensic applications such as on-site identification of crime scene samples, identification of victims after a mass disaster, or for border control to fight human trafficking. Indeed, these applications will require further development of the extraction methods to ensure sufficient DNA is available for sequencing. With the upcoming early release of the Voltrax (an automated library preparation device) and the Zumbador project (a complete device for DNA extraction and sample preparation), these portable sample preparation techniques might soon be available. Furthermore, ONT recently announced the development of SmigION, a nanopore-based sequencer that will be plugged into a cellphone (Yong, 2016). With this invention, MinION sketching can eventually promote a range of futuristic Internet of (living) Things applications that will use DNA as a means for biometric authentication.

MinION sketching provides a rapid method for cell authentication and sample re-identification. We developed and implemented a Bayesian method that allows matching error-prone MinION reads to sparse matching files from a database. We showed the robust matching and specificity of DNA sample re-identification using 91-250 SNPs. This creates the opportunity for large-scale implementation in research labs, clinical settings and forensics. Databases for cell line authentication can be easily constructed using available online genomic data. To kick-start the initiative, we provide the 1099 cancer genome reference files generated by the CCLE in a format compatible with our pipeline.

## Methods

### The Bayesian matching algorithm

The matching algorithm uses a Bayesian framework to evaluate the posterior probability of a match. Let *x*_*i*_ ∈ {*Y*, *N*} be a random variable that either indicates whether the MinION sketch directly matches a known person (*x*_*i*_ = *Y*), or does not match (*x*_*i*_ = *N*) with respect to the *i*-th individual in the database. Let *D*_*k*_ be the observed MinION data for the *k*-th bi-allelic marker, with *D*_*i*_ ∈ {*A*, *B*}, where *A* and *B* denote the two alleles; and Let ***D*** = (*D*_*i*_, *D*_*i*_,…, *D*_*i*_) denote the observation for *n* bi-allelic markers.

The posterior probability of the matching outcome for the *i*-th sample is:

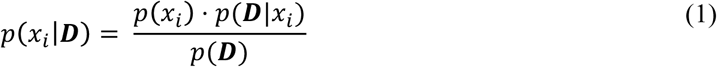

where *p* (*x*_*i*_) is the prior probability for the matching status of *i*-th sample and is specified by the user.

The likelihood is approximated using the following equation:

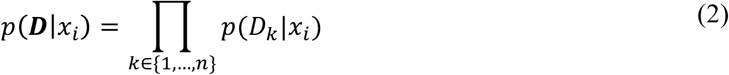

The likelihood of an <u>exact match</u> given the data of the *k*-th marker, *p*(*D*_*k*_|*x*_*i*_ = *Y*), is given by the following matrix:

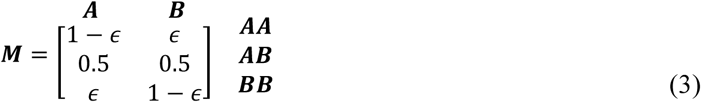

where the rows denote the genotype of the *i*-th sample for the *k*-th marker as observed in the DNA database, the columns correspond to the observed genotype in the MinION data, and *ϵ* denotes the error rate assuming symmetry in confusing allele *A* for allele *B* and *vice versa*. *p (D*_*k*_\*x*_*i*_ = *Y*) corresponds to a specific row of ***M*** based on the observed genotype of a sample in the database. For example, if the genotype of the database sample is *AA*, then *p (D*_*k*_ = *A x*_*i*_ = *Y*) = 1 − *ϵ* and *p (D*_*k*_ = *B x*_*i*_ = *Y*) = *ϵ*.

The likelihood of a <u>mismatch</u> given the data of the *k-*th marker, *p*(*D*_*k*_|*x*_*i*_ = *N*), basically corresponds to observing the allele *D*_*k*_ in a random person from the population. This probability is the sum of two processes: (i) the random person has the same allele as *D*_*k*_ and the observation is errorless or (ii) the random person does not have the same allele as *D*_*k*_ but a sequencing error flipped the observed allele. Therefore:

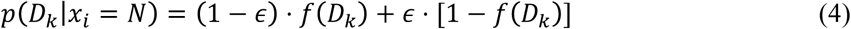

where *f*(*D*_*k*_) denotes the frequency of the observed allele in the population.

Finally, the evidence, *p* (***D***) is given by:

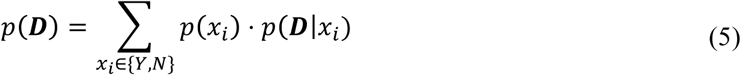

### DNA samples for sequencing

We purchased the genomic DNA sample for the 1000 Genomes individual NA12890 from the Coriell Institute. The THP1 cell line (ECACC: 88081201 sigma) was used from the lab recourses. YE001 and SZ001 were derived from the corresponding authors (Y.E. and S.Z.) and JP001 using a saliva collection kit or cheek-swabs. DNA preparation of 2D libraries was done as in Zaaijer & Erlich, 2016 (see also: supplemental materials). Rapid library and barcoding for MinION sequencing: according to manufacturers directions (see also: supplemental materials).

### DNA samples as a reference database

YE001, JP001 and three HapMap samples (NA12890, NA12977, NA12879) are publicly available reference files. The 1099 cancer cell line files were downloaded, base-called using Birdseed and converted into 23andMe file format. The 31,000 DTC genomes were available from two sources: (i) 1446 DTC genomes were downloaded from the public website OpenSNP.org and (ii) 29,554 genomes were collected using DNA.Land, an online website (<u>https://dna.land</u>). The website procedures were approved by our IRB. Based on current consent, this set of 29,554 genomes cannot be shared. All experiments with this collection were done using an automatic algorithm on a secure server without access to the explicit identifiers of the samples (e.g. names or contact information) (further information in Supplemental Materials).

### MinION sketching

The MinION was run according to the instructions of the manufacturer. We used Poretools (Loman & Quinlan 2014) to extract the FASTQ data and time stamps from the local files, followed by alignment using bwa-mem (Li, 2013). Only SNPs present in dbSNP build-138 with an allele frequency between 1-99% were selected. The Bayesian model was integrated in a Python script, in order to match between the MinION data and each entry in the database. As a default setting, we used a prior probability of 10^-5^ for exact matching. All code is publicly available on github at github.com/TeamErlich/personal-identification-pipeline.

## Declarations

### Ethics approval

All individuals (YE001, JP001, SZ001) declare they fully consented to participate in the study.

### Availability of the data

The code for our method is available on <u>https://github.com/TeamErlich/personal-identification-pipeline</u>. We also include a reference database for the CCLE cell line repository for fast re-identification.

### Author contributions

S.Z. and Y.E. designed the experiments and wrote the manuscript. S.Z. conducted the sequencing experiments, developed the portable sketching method, and analyzed the data. R.P., D.S. and Y.E. devised the Bayesian algorithms. A.G., R.P., D.S., and Y.E. coded the algorithm.

## Acknowledgements

We thank Eleazar Eskin for useful comments on the model. We thank Neville Sanjana for providing cell lines and for discussions. We thank D. Zielinski and T. Willems for useful comments, W. Stephenson and K. Pandit from New York Genome Center’s Innovation lab for technical assistance and A. Wolman for providing the THP1 cell line. We thank M. Micorescu from Oxford Nanopore Technologies for useful discussions, and the Columbia Ubiquitous Genomics class 2015 for data generation.

## Supplemental material

Table of contents:

- Supplemental figure 1 -3
- Supplemental Tables 1-5
- Supplemental experimental procedures

**Supplemental figure 1.**
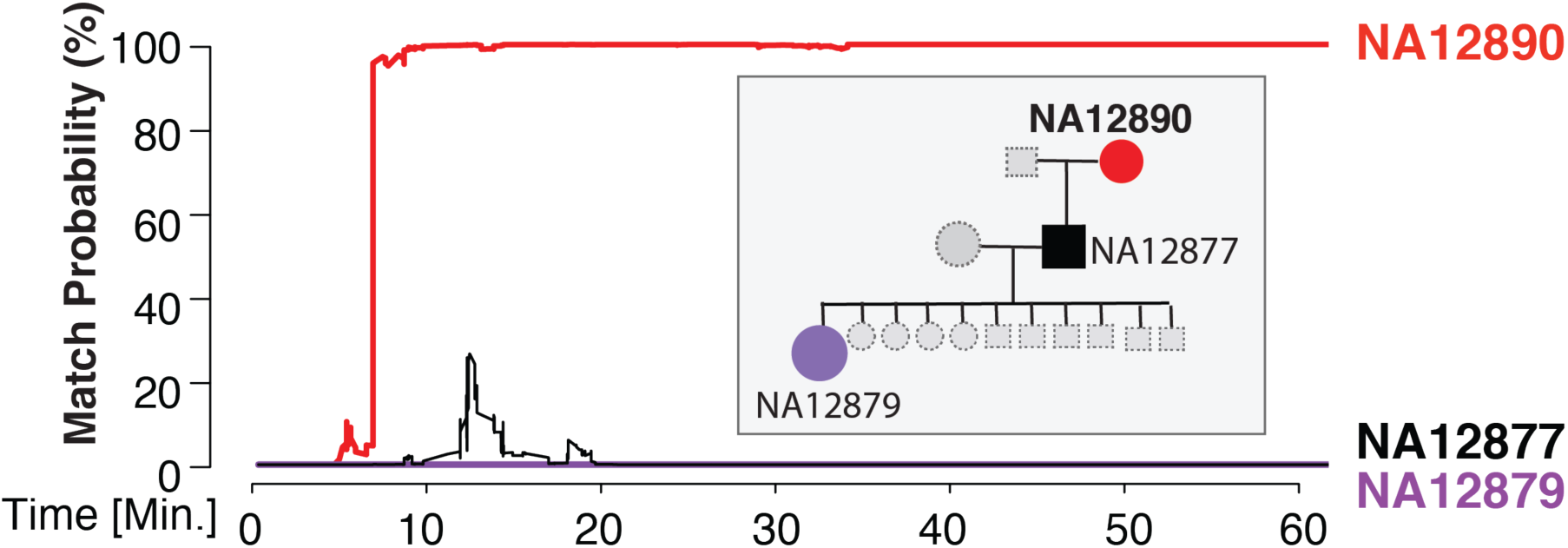
Results of sketching NA12890. (a) The pedigree of 1000Genomes sample NA12890 (b) The posterior probability for an exact match between the sketch of NA12890 and her genome (red), her son’s genome (black), and her granddaughter’s genome (purple) as a function of sketching time.

**Supplemental figure 2.**
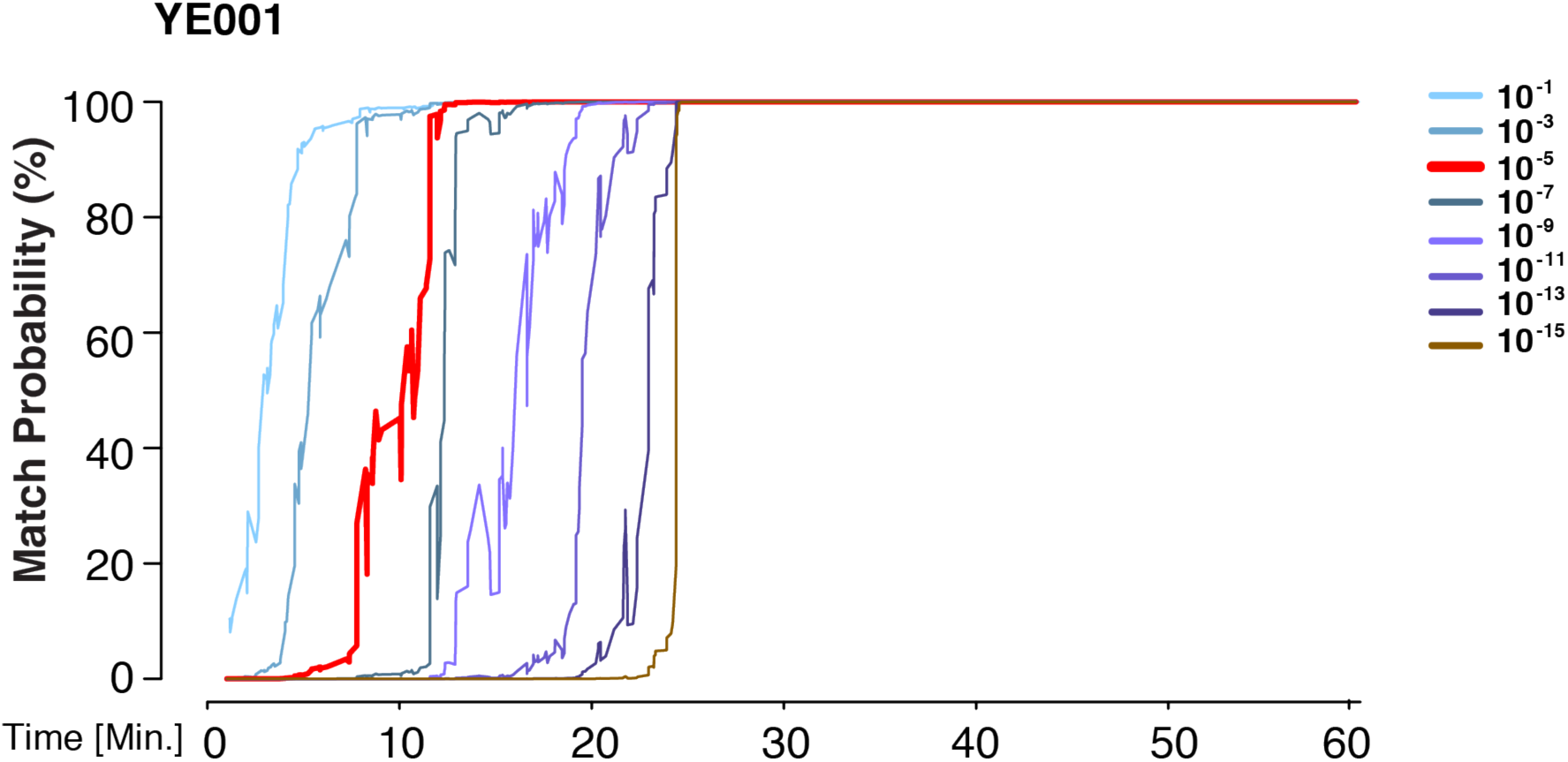
Prior representing a database larger then the world population still allows identification power. The match probability is inferred by comparing a MinION sketch of YE001 to their reference file as a function of the MinION sketching time. The prior probability for a match was modified as indicated.

**Supplemental figure 3.**
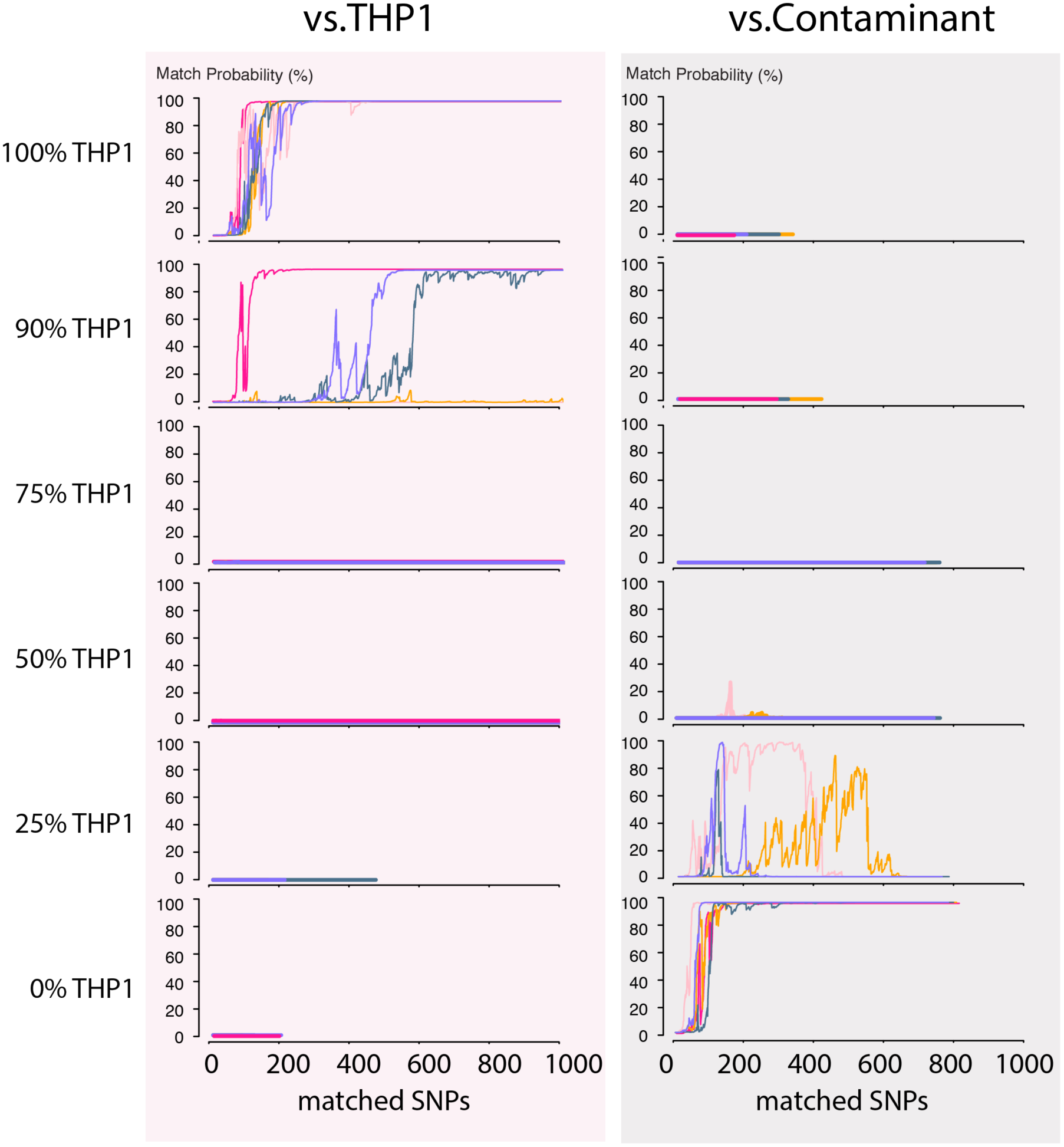
Contamination simulations. Random reads from a run with THP1 cells are mixed in the indicated proportions and shuffled. This simulated MinION sketch is matched against the THP1 reference file, and the contaminant reference file. This process is repeated five times for each simulated contamination (pink, light-pink, purple, green and yellow lines). The match probability is here a function of the number of SNPs used in the Bayesian.

## Supplemental experimental procedures

### DNA preparation for 2D sequencing

Genomic DNA from NA12890 and YE001 (Table S1; exp1, exp2 respectively) were prepared for 2D MinION libraries (SQK-MAP006 ONT) as described by Zaaijer et al., 2016. 2D libraries are double stranded DNA fragments with a ligated hairpin loop and adaptors containing a tether and motor protein necessary for MinION sequencing, these are run on the R7 flow-cells. DNA samples from SZ001, JP001 and the THP1 cell line were prepared using the SQK-NSK007 (Table S1; exp3, exp 4, exp 5) and run on R9 flowcells.

### Rapid library preparation in the lab

Samples (Table S1, exp6) were collected by cheek swap (Catch-All™ Sample Collection Swab Epicentre QEC89100) scraping ∼30 sec both sides of the cheek. Cells were recovered in 200ul PBS. After addition of 20μl Proteinase K and 200μl lysis buffer (DNeasy blood & tissue kit, Qiagen, #69504) the sample was incubated at 56^o^C for 10 minutes. The sample is then applied to the column, spun 1 minute, followed by two wash steps with AW1 and AW2 respectively. Next, 20 μl elution buffer was applied and the column was spun for 1 minute on a regular benchtop centrifuge at max speed. Recovery of the DNA sample in 20μl resulted in an average yield of ∼3-5ng/ μl.

We used the SQK-RAD001 kit to prepare the DNA library. FRM (2.5μl, ONT) was added to the DNA sample (20μl) and incubated for 1 min at 30^o^C. Then, 1μl RAD (ONT) plus 0.2μl ligase was added and the mixture was incubated for 10 minutes.

The R9 flowcell was prepared by applying two times 500ul priming mix (RBF 1x). The library was then added to the flowcell without a purification step.

### Barcoding

The barcoding protocol was executed according to manufacturer’s instructions for native barcoding kit I (EXP-NBD002) in conjunction with Nanopore Sequencing kit (SQK-NSK007) with some modifications (Table S1, exp. 5, exp. 4). In brief; 1.5 ug DNA was used for each sample as starting material and vigorously vortexed for a minute. The DNA sample was end-repaired and dA-tailed using the NEBNext Ultra II End Repair/dA-tailing Module (5 min 20^o^C, and 5 min 65^o^C). After an AMPure purification, the DNA fragments were subject to ligation using Blunt/TA Ligase Master Mix (NEB M0367S) for 5 minutes at 20^o^C and then 5 minutes at 65^o^C. The sample was then purified using AMPure magnetic beads and the DNA was eluted off the beads using 31μl nuclease free water (NFW). The NB01 and NB02 barcode was ligated to the fragments of each sample with Blunt/TA ligase mix (NEB) and incubated for 15 minutes. After an AMPure purification step, the two samples are pooled. Next we ligated the adaptor (BAM) and hairpin (BHP) to the barcoded DNA fragments using NEB quick ligase (NEB) for 20 minutes at room-temperature (22^o^C). The HTP (ONT) was added and incubated for another 10 minutes. The 50 ul MyOne C1 beads were prepared in the incubation step, which tethers the hairpin and ligated DNA fragments. The DNA library was eluted off the beads by ELB (ONT) at 37^o^C for 10 minutes and was applied to the flow cell.

### MinION sketching

To start a MinION run, we primed the flowcells according to the manufacturer’s protocol. We started MinKnow (protocol “MAP_48Hr_Sequencing_Run_SQK_MAP006” for R7 and “NC_48hr_Sequencing_Run_FLO-MIN104” for R9), uploaded the collected reads to Metrichor (a cloud-based program that base-called the reads), and stored them on our computer.

We used Poretools [(Loman & Quinlan 2014)] to extract the FASTQ data and time stamps from the local files. Only reads with an average base quality greater than 9 were used for the downstream analysis. Next, we aligned the files to hg19 using bwa-mem (v0.7.14)(Li 2013) using the command “bwa mem –V –x ont2d –t 4”. Reads with multiple alignments were not considered for further analysis.

To extract variants, we used a custom script to retain nucleotides from the MinION output that overlap known positions of bi-allelic SNPs from dbSNP build-138 with an allele frequency between 1-99%. To minimize the effects of sequencing error, we considered only MinION read bases that matched the common SNP alleles in dbSNP. For example, if at position chr1:10,000 the MinION reported “A” and dbSNP reported a variant “C/G”, then we treated this position as a sequencing error. The R7 chemistry run with NA12890 generated 4920 variants after one hour of MinION sequencing, of which 7.7% were rejected after filtering for common SNPs. Intersecting these with the reference file and analyzing the true error from the matched SNPs resulted in 8.9% mismatches. This contrasts with the R9 chemistry, which only resulted in 2% true mismatches **(Table S3-5**).

The Bayesian model was integrated in a Python script, in order to match between the MinION data and each entry in the database. To accelerate the search, we implemented the following procedure: (i) if the posterior probability drops below 10^-9^, the script concludes that the database entry does not match and moves to the next entry (ii) the script uses only up to one hour of data to determine the posterior of a sample.

As a default setting, we used a prior probability of 10^-5^ for exact matching. The only exception was Figure supplement 2 (YE001), where we employed a range of prior probabilities. As a default setting, we used the computed error rate from each read as the *ϵ* in our Bayesian.

All code is publicly available on github at github.com/TeamErlich/personal-identification-pipeline.

### Simulations

For the simulations we took reads from exp. 4 and 5 (Table S1). The total number of reads was set to 3000 and a random number of reads that represents the percentage proportion were selected. For example, for 50% contamination we took 1500 random reads from experiment 4 and 1500 random reads from experiment 5. These were pooled together and again shuffled to simulate a mix. This process was repeated five times for each contamination fraction. The resulting pooled file was processed using our pipeline and matched to the reference file of the corresponding MinION sketch (either THP1, or JP001).

**Table S1.**
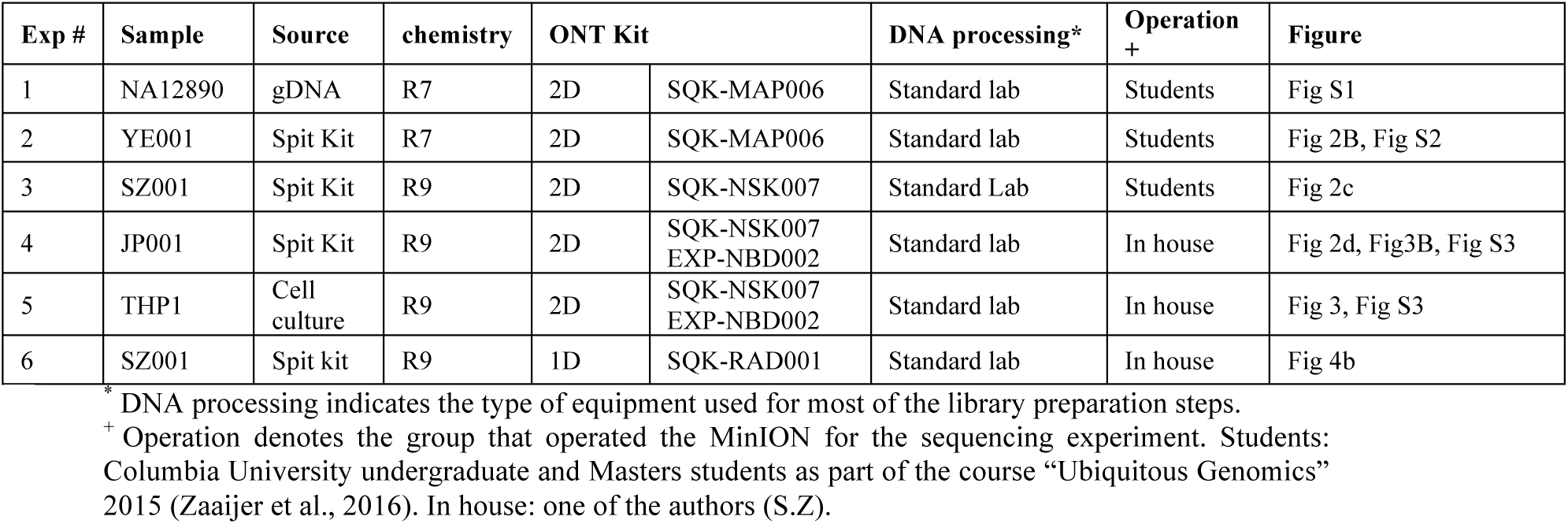
Experimental Summary.

| Exp # | Sample | Source | chemistry | ONT Kit |  | DNA processing* | Operation + | Figure |
| --- | --- | --- | --- | --- | --- | --- | --- | --- |
| 1 | NA12890 | gDNA | R7 | 2D | SQK-MAP006 | Standard lab | Students | Fig S1 |
| 2 | YE001 | Spit Kit | R7 | 2D | SQK-MAP006 | Standard lab | Students | Fig 2B, Fig S2 |
| 3 | SZ001 | Spit Kit | R9 | 2D | SQK-NSK007 | Standard Lab | Students | Fig 2c |
| 4 | JP001 | Spit Kit | R9 | 2D | SQK-NSK007<br>EXP-NBD002 | Standard lab | In house | Fig 2d, Fig3B, Fig S3 |
| 5 | THP1 | Cell culture | R9 | 2D | SQK-NSK007<br>EXP-NBD002 | Standard lab | In house | Fig 3, Fig S3 |
| 6 | SZ001 | Spit kit | R9 | 1D | SQK-RAD001 | Standard lab | In house | Fig 4b |

\* DNA processing indicates the type of equipment used for most of the library preparation steps.
+ Operation denotes the group that operated the MinION for the sequencing experiment. Students: Columbia University undergraduate and Masters students as part of the course “Ubiquitous Genomics” 2015 (Zaaijer et al., 2016). In house: one of the authors (S.Z).

**Supplemental Table S2.**

|  | NA12890<br>2D | YE001<br>2D |
| --- | --- | --- |
|  | **Sequencing yield** |  |
| Passed bases (#) | 17,675,127 | 48,451,196 |
| Passed reads (#) | 2,272 | 10,067 |
| Read length average (bp) | 7,779 | 4,812 |
| Unique aligned reads (#) | 1,451 | 7,808 |
| Aligned bases (#) | 27,810 | 112,988 |
| Avg. read error rate (%) | 9.6 | 7.4% |
|  | **Matching details** |  |
| **#SNPs to positive identification\*** | **195** | **110** |
| Match homozygous genotype | 54 | 76 |
| Homozygous mismatch | 10 | 2 |
| Match heterozygous genotype | 131 | 7 |
| Time to positive identification (min.) | 13min | 13min |

\*positive identification was defined as 99.9% for 2D experiments

**Supplemental Table S3.**

|  | SZ001<br>2D | JP001<br>2D |
| --- | --- | --- |
|  | **Sequencing yield** |  |
| Passed bases (#) | 33,216,820 | 21,369,107 |
| Passed reads (#) | 8,610 | 7,425 |
| Read length average (bp) | 3,857 | 2,878 |
| Unique aligned reads (#) | 6,127 | 5,783 |
| Aligned bases (#) | 98,504 | 67,402 |
| Avg. read error rate (%) | 3.8 | 3.4 |
|  | **Matching details** |  |
| **#SNPs to positive identification\*** | **98** | **134** |
| Match homozygous genotype | 66 | 88 |
| Homozygous mismatch | 3 | 4 |
| Match heterozygous genotype | 29 | 42 |
| Time to positive identification (min.) | 11.4min | 4.7min |

\*positive identification was defined as 99.9% for 2D experiments

**Supplemental Table S4.**

|  | THP1<br>pure | THP1<br>contaminated |
| --- | --- | --- |
| **Sequencing yield** |  |  |
| Passed bases (#) | 11,721,501 | 31,283,238 |
| Passed reads (#) | 3,823 | 9,555 |
| Read length average (bp) | 3,066 | 3,274 |
| Unique aligned reads (#) | 3,594 | 8,991 |
| Aligned bases (#) | 38,135 | 98,705 |
| Avg. read error rate (%) | 5.24 | 5.20 |
| **Matching details** |  |  |
| #SNPs to positive identification* | 91 |  |
| Match homozygous genotype | 72 |  |
| Homozygous mismatch | 1 |  |
| Match heterozygous genotype | 18 |  |
| Time to positive identification (min.) | 3min |  |

\*positive identification was defined as 99.9% for 2D experiments

**Supplemental Table S5.**
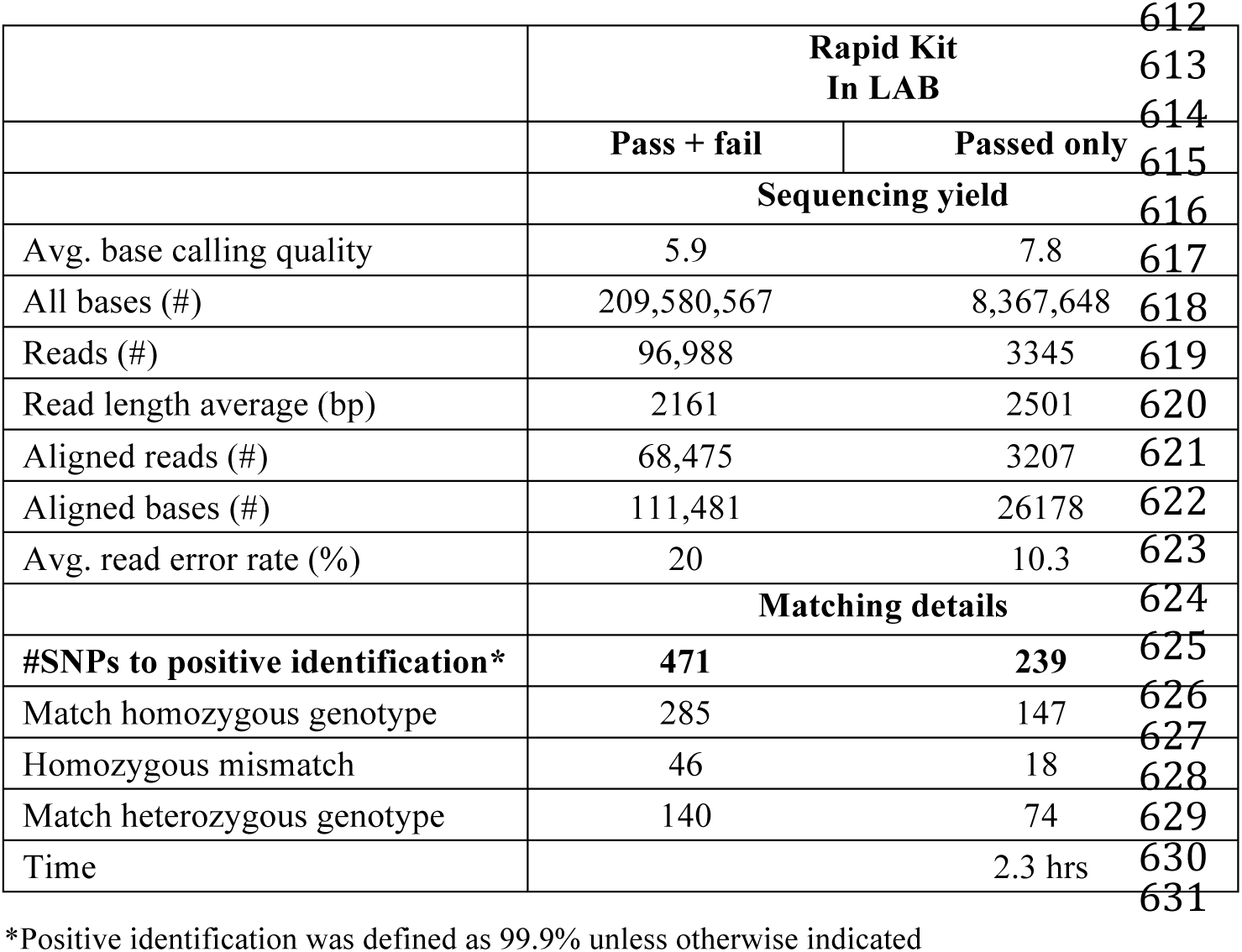

|  | **Rapid Kit**<br>**In LAB** |  |
| --- | --- | --- |
|  | Pass + fail | Passed only |
| **Sequencing yield** |  |  |
| Avg. base calling quality | 5.9 | 7.8 |
| All bases (#) | 209,580,567 | 8,367,648 |
| Reads (#) | 96,988 | 3345 |
| Read length average (bp) | 2161 | 2501 |
| Aligned reads (#) | 68,475 | 3207 |
| Aligned bases (#) | 111,481 | 26178 |
| Avg. read error rate (%) | 20 | 10.3 |
| **Matching details** |  |  |
| #SNPs to positive identification* | 471 | 239 |
| Match homozygous genotype | 285 | 147 |
| Homozygous mismatch | 46 | 18 |
| Match heterozygous genotype | 140 | 74 |
| Time |  | 2.3 hrs |

\*Positive identification was defined as 99.9% unless otherwise indicated

